# Symmetry-Based Center and Rotation Refinement for Fiber Diffraction Patterns

**DOI:** 10.64898/2026.08.22.746299

**Authors:** Irina Klein, Gady Agam, Thomas Irving

## Abstract

X-ray fiber diffraction patterns exhibit four-fold symmetry that can be exploited, through folding and averaging, to improve signal-to-noise ratio. Accurate folding requires a precise subpixel estimate of the symmetry center and precise orientation of the meridional pattern axis to the fiber axis: small center or angular errors blur diffraction features, reduce layer-line sharpness, and introduce errors in spacing measurements. A pixel-level estimate is often too imprecise for this purpose, and detector gaps further complicate the alignment objective. We formulate the masked quadrant-folding problem, define a four-quadrant symmetry loss that consistently excludes invalid pixels, and evaluate several refinement strategies: hierarchical coarse-to-fine grid search; ECC-based rigid registration with global center/orientation correction fitting; ECC registration followed by local gradient refinement; and a hybrid that appends a local grid search on a cropped pattern. Direct gradient optimization from the rough QF alignment was found to be unreliable. Grid search provides a robust, interpretable baseline that directly minimizes the folding objective but is substantially slower than registration; ECC gives a fast near-correct alignment, and the hybrid closes the accuracy gap to brute-force search at a fraction of its runtime. On real datasets with calibration data, applying a calibration center with optimized rotation is effectively optimal. The hybrid center-refinement method has been integrated into the MuscleX package.

## 1 Introduction

X-ray fiber diffraction images contain structured intensity patterns that reflect the organization of the underlying molecular or material system. In many experimental settings the diffraction pattern is expected to exhibit four-fold symmetry with respect to the meridional and equatorial axes. This symmetry can be exploited by folding the image: the four quadrants are reflected into a common coordinate frame and corresponding pixels are averaged. When the symmetry model is valid, folding improves signal-to-noise ratio, reduces redundancy, and provides a more stable representation for downstream measurements such as layer-line tracking, peak fitting, and temporal state analysis.

Accurate folding depends critically on the estimated symmetry center and orientation of the pattern. If the assumed center is displaced or the assumed symmetry axes are rotated relative to the true diffraction pattern, corresponding pixels from different quadrants will not sample the same physical feature. Even small center or angular errors can blur sharp diffraction structures, reduce layer-line contrast, distort intensity measurements, and introduce errors in spacing measurements. Alignment refinement is therefore an important preprocessing step before quadrant folding.

The problem is further complicated by the presence of detector gaps and other invalid image regions. These regions should not contribute to folding, symmetry evaluation, or alignment refinement. If gap pixels or invalid regions are included in the objective, the refinement procedure may partially align detector artifacts rather than diffraction features. A useful alignment method must therefore combine the image geometry, the symmetry model, and the binary validity mask in a consistent way.

This report formulates the masked quadrant-folding problem and evaluates practical methods for refining the symmetry center and orientation for muscle fiber diffraction patterns. The methods assume that a rough alignment is already available and focus on estimating a small correction to the center and orientation. A masked four-quadrant symmetry loss is defined, and several refinement strategies are considered: direct symmetry-loss optimization, hierarchical coarse-to-fine grid search, rigid registration of quadrants, local gradient refinement initialized from registration, and a hybrid approach combining the strengths of the previous methods. The results support a hybrid strategy. Hierarchical grid search is robust and interpretable because it directly evaluates the final folding objective, but it is relatively slow. It can, however, be sped up by using a smaller image with cropping to the central region, where the majority of the diffraction signal is located. ECC-based rigid registration is much faster and provides a near-correct alignment by estimating relative quadrant motions and fitting them to a physically valid global center/orientation correction. Direct gradient optimization from the original rough alignment was not reliable, but gradient refinement initialized from the ECC result improved on registration alone. Applying a fine-resolution local search on top of the ECC registration-and-gradient refinement further improved the result. The recommended pipeline is therefore

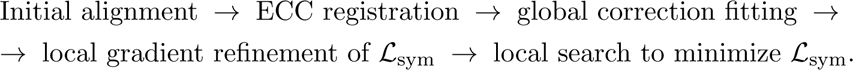

### Contributions

- A *masked* four-quadrant symmetry loss that consistently excludes detector gaps across extraction, folding, registration, and scores the quality of quadrant folding.
- Four refinement strategies and a hybrid pipeline (“Reg+Grad+Search”) that gives the best loss reduction on most datasets at lower runtime than that of brute-force search, enabling calibration-quality center without a calibration image.
- Mask-aware ECC extensions (erosion, re-seeding, dynamic reference selection) and a coarse-to-fine variant, evaluated with ablations.
- A controlled comparison against a per-dataset calibration baseline showing calibration-with-optimized-rotation is effectively optimal.
- An independent reliability study of pyFAI calibration showing that calibration image quality governs the recovered center.

Figure 1 illustrates the task and its payoff on a representative muscle fiber diffraction pattern: the original pattern on the left is off-center and rotated relative to its symmetry axes, the rough MuscleX QF estimate brings it close, and the refinement methods (grid search and ECC registration) recover a sharper, better-centered alignment with a lower symmetry loss.

**Figure 1:**
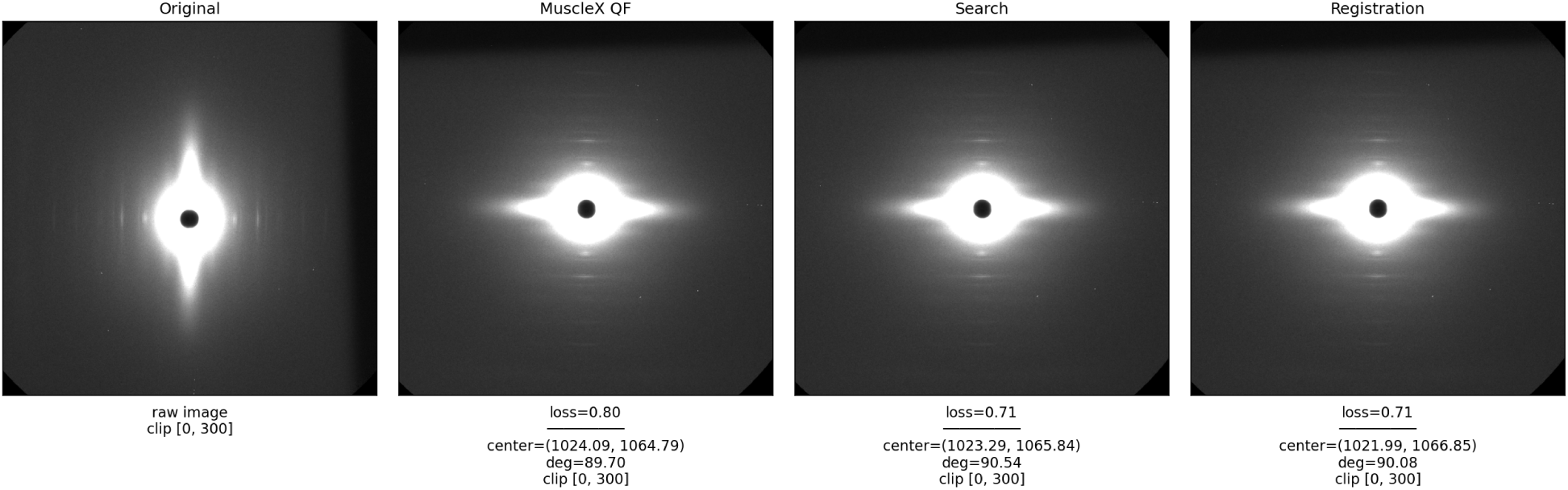
The refinement task on a representative pattern. From left to right: original image; the same image aligned using the MuscleX QF estimate; and the alignments recovered by brute-force Search and by ECC registration. Captions report the masked symmetry loss *L^′^*_Sym_ together with the estimated center and rotation; both refinements lower the loss relative to QF.

The remainder of the report is organized as follows. Section 2 positions the method against detector calibration, the MuscleX quadrant-fold baseline, image registration, and symmetry-based alignment. Section 3 formulates masked quadrant folding and defines the masked symmetry loss. Section 4 presents the four refinement strategies, the hybrid pipeline, and the mask-aware ECC extensions. Section 5 describes the datasets, baselines, metrics, and implementation. Section 6 reports the synthetic validation, the multi-dataset evaluation, the configuration studies, the calibration comparison, and the comparison of MuscleX calibrated center to pyFAI. Sections 7–9 discuss the findings and state the limitations.

## 2 Related Work

Aligning a diffraction pattern before quadrant folding draws on three bodies of work: detector calibration and beam-center estimation, image registration, and symmetry-based alignment. We review each below and position the present method, which refines an already-rough alignment by directly minimizing a masked folding loss.

### 2.1 Detector calibration and beam-center estimation

When a dedicated calibration exposure of a powder calibrant standard such as silver behenate (AgBh) is available, the detector geometry and therefore the beam center can be recovered by fitting the known ring radii of the powder rings [7]. Established tools implement this azimuthal-geometry refinement, including FIT2D [6] and pyFAI [1, 10], the last of which we use as the calibration reference in Section 6. These methods are accurate but require a separate calibration image and assume the presence of multiple complete diffraction rings. The reliability of pyFAI results in the presence of detector gaps and sparse rings is a question we examine empirically in Section 6.

### 2.2 Fiber-diffraction analysis and the MuscleX quadrant-fold baseline

MuscleX is an open-source suite for muscle fiber diffraction analysis [9], and its quadrant-fold (QF) routine supplies both the per-image initial alignment (the “QF baseline” used in this work) and the normalized fold standard deviation used as the symmetry loss. MuscleX determines the center in one of two ways: from a calibration image when one is supplied, or directly from each diffraction image otherwise, which is the default for the datasets refined here.

#### With a calibration image

The user selects at least five points along an AgBh [7] calibration ring. Because manual clicks do not fall exactly on the intensity ridge, each point is then refined: the image is sampled along the radial direction connecting the initial center to the point over a ±30-pixel window using bilinear interpolation, and the point is relocated to the local intensity maximum. An ellipse is refit to the refined points to update the center, with the ring radius taken as the mean of the two semi-axes. The estimate is finalized by a contrast-based optimization over the center coordinates and radius jointly. The objective samples intensity in a narrow signal band on the ring and in a pair of background bands around the ring, and returns the band contrast (mean signal on the ring minus a weighted mean background off the ring). This objective forces the fit to be on the brightest part of the ring. The objective is maximized by differential evolution [14] within ±100 pixels of the center estimate and ±10% of the radius estimate, followed by local L-BFGS-B polishing [4], yielding the calibrated center and ring radius. The radius, combined with the standard’s known *d*-spacing, fixes the detector scale.

#### Without a calibration image: automatic center determination

MuscleX estimates the diffraction center through a cascade of methods, returning the first that yields a result consistent with an internal check. The input image is converted to 8-bit and smoothed with a 5 × 5 Gaussian kernel. An initial estimate is obtained by thresholding the image to retain the brightest 0.5% of pixels, applying a morphological opening with a 50-pixel elliptical structuring element to suppress isolated noise, extracting contours, and fitting an ellipse to the longest contour; the ellipse centroid serves as the provisional center. This estimate is then refined using the diffraction reflections: the image is thresholded more aggressively (retaining the brightest 0.015% of pixels) to isolate discrete reflection spots, ellipses are fitted to the six largest resulting contours, and the pair of reflections with the most similar integrated areas is identified under the assumption that symmetric reflections on each side of the true center are of comparable size. When no contour-based estimate can be formed, the method falls back successively to the intensity-weighted centroid computed from image moments, to a Hough circle fit, and lastly to the geometric center of the image.

#### Rotation determination

The rotation angle of the diffraction pattern is computed by combining an ellipse-based estimate with azimuthal integration. As in the center step, the image is converted to 8-bit, Gaussian-smoothed, and thresholded, and an ellipse is fitted to the longest contour. The ellipse orientation provides an initial angle. The image is then azimuthally integrated about the determined center into a two-dimensional intensity map of radius versus angle. The map is truncated to the inner third of the radial range, where the signal dominates, and summed over radius to yield an intensity-versus-angle profile. The orientation is taken as the angle maximizing the summed intensity of a diametrically opposed bin pair. A second, finer integration over a ±5*^◦^* window around this angle and its complement refines the estimate to sub-degree resolution. If the integration-derived angle and the ellipse-derived angle differ by more than 20*^◦^*, the ellipse angle is returned in preference; otherwise the integration angle, folded to the range [−90*^◦^,* 90*^◦^*], is returned.

No-calibration center and rotation estimators are both heuristic and per-image: they rely on thresholding and contour fitting that are sensitive to noise, detector gaps, and weak reflections, and they do not optimize the eventual folding objective. This motivates a refinement step that directly minimizes the folding loss.

### 2.3 Image registration

Image registration recovers a geometric transform aligning two images and is broadly split into feature-based and intensity-based methods [16]. We use the intensity-based enhanced correlation coefficient (ECC) criterion of Evangelidis and Psarakis [5], which estimates Euclidean motion by maximizing a normalized correlation that robustly handles variations in lighting and contrast, available as findTransformECC in OpenCV [3]. We extend standard ECC with mask erosion, re-seeding, and dynamic reference selection so that detector gaps do not corrupt the alignment (Section 4).

### 2.4 Symmetry-based alignment and optimization

Estimating a center and axes of symmetry from image content is a long-studied problem in computer vision [12]; our objective specializes it to mirror (reflection) symmetry across two orthogonal axes under a masked folding average. To optimize this objective we combine a coarse-to-fine search — a classical strategy for escaping poor local minima in alignment problems [2] — with local refinement using the bound-constrained L-BFGS-B optimizer [4], optionally under a robust Huber penalty [8] to limit the influence of residual asymmetries. The supporting numerical machinery is provided by SciPy [15] and OpenCV [3].

### 2.5 Positioning of the proposed refinement method

Unlike calibration-based methods, our refinement needs no separate calibration exposure: it operates on each diffraction image and corrects the existing QF alignment. Unlike the heuristic QF estimators, it optimizes the actual folding objective. And unlike off-the-shelf registration, it is mask-aware throughout and projects the per-quadrant transforms onto a single physically valid center/orientation correction. We further benchmark against a calibration baseline. We also separately audit the reliability of the pyFAI calibration as a comparison to the MuscleX calibration approach.

## 3 Problem Formulation

### 3.1 Quadrant symmetry model and masked folding

Consider a sequence of two-dimensional diffraction X-ray images in which the underlying diffraction pattern is expected to exhibit approximate quadrant symmetry. The goal is to fold each image by reflecting its four quadrants into a common coordinate frame and averaging corresponding pixels. Accurate folding requires an accurate estimate of both the symmetry center and the orientation of the symmetry axes; even small center or angular errors blur sharp features, distort layer-line intensities, or introduce artifacts into the folded image.

Let *I* : Ω ⊂ ℝ^2^ → ℝ denote the diffraction image intensity, and let *M* : Ω → {0, 1} denote a binary invalid-pixel mask, where *M* (***x***) = 1 indicates that the pixel should be ignored and *M* (***x***) = 0 indicates that the pixel is valid. The mask accounts for detector gaps, dead pixels, beamstop regions, saturated regions, or other image locations that should not participate in alignment, symmetry evaluation, or averaging.

The rough alignment of an image is described by a center ***c*** = (*c_x_, c_y_*)*^T^* and an orientation angle *θ*. The center defines the intersection of the two symmetry axes, and *θ* defines the orientation of those axes relative to the image coordinate system. The alignment parameter vector is ***p*** = (*c_x_, c_y_, θ*). Given an initial rough alignment ***p***_0_ = (*c_x_*_0_*, c_y_*_0_*, θ*_0_), the objective is to estimate a small correction Δ***p*** = (Δ*c_x_,* Δ*c_y_,* Δ*θ*) such that the refined alignment ***p****^∗^* = ***p***_0_ + Δ***p*** produces improved agreement among corresponding pixels across the four reflected quadrants, while excluding masked pixels.

The central difficulty is that the three alignment parameters are coupled. A center error can appear as relative translations between reflected quadrants, while an orientation error can also produce apparent translations or local rotations. In addition, detector masks, interpolation, sharp diffraction features, repeated layer-line structures, and approximate rather than perfect physical symmetry can make the objective function non-smooth or locally ambiguous. These factors make direct optimization from a rough initial estimate unreliable in some cases.

### 3.2 Canonical folded coordinates

Let ***u*** = (*u_x_, u_y_*)*^T^* denote a point in a canonical folded-quadrant coordinate system. This coordinate system represents the common quadrant into which all four original image quadrants are reflected before averaging. In this frame, corresponding physical diffraction features should appear at the same coordinate ***u*** when the center and orientation are correctly estimated.

The four reflected quadrants are indexed by Q = {++, +−, −+, −−}. Each quadrant is represented by a sign matrix that reflects the canonical coordinate across one or both symmetry axes:

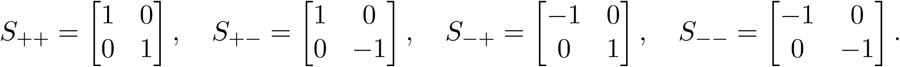

The orientation of the symmetry axes in the original image is represented by the rotation matrix

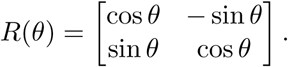

For a given alignment parameter vector ***p*** = (*c_x_, c_y_, θ*) with center ***c*** = (*c_x_, c_y_*)*^T^*, the mapping from a canonical folded-quadrant coordinate ***u*** to the original image coordinate in quadrant *q* is

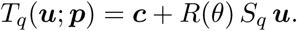

The corresponding sampled quadrant image is *Q_q_*(***u***; ***p***) = *I*(*T_q_*(***u***; ***p***)), and the corresponding validity weight is *W_q_*(***u***; ***p***) = 1 − *M* (*T_q_*(***u***; ***p***)), so that *W_q_* = 1 indicates a valid sample and *W_q_* = 0 indicates a sample to be ignored. In implementation the image intensity *I* is sampled using interpolation, while the binary mask *M* is sampled or thresholded so that invalid detector regions do not contribute to folding or loss evaluation.

### 3.3 Interpretation of corresponding pixels

In a perfectly centered and oriented image, the four quadrant samples at the same canonical coordinate ***u*** correspond to the same physical diffraction feature. For example, in a 100 × 100 image with ideal horizontal and vertical symmetry axes aligned with the image grid, the pixels (0, 0), (0, 99), (99, 0), and (99, 99) are corresponding corner pixels under quadrant reflection. More generally, corresponding pixels are obtained by reflecting points across the two symmetry axes defined by the center ***c*** and orientation *θ*.

This model represents reflection-based quadrant (mirror) symmetry, not fourfold rotational symmetry. The matrices *S_q_* encode sign changes across the two symmetry axes, and the rotation *R*(*θ*) places those axes in the original image coordinate system. This distinction matters for diffraction patterns in which the expected symmetry is mirror symmetry across the meridional and equatorial axes rather than invariance under 90*^◦^* rotations. Under the correct alignment, the four quantities *Q_q_*(***u***; ***p***) should agree up to measurement noise, detector effects, and physical deviations from perfect symmetry.

### 3.4 Masked folding

For a given alignment ***p***, the folded intensity at canonical coordinate ***u*** is the masked average of the valid quadrant samples:

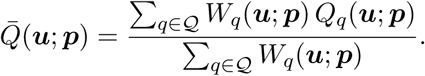

This expression is evaluated only when at least one valid quadrant sample is available; the valid-sample count is *N* (***u***; ***p***) = ∑*_q∈Q_ W_q_*(***u***; ***p***), and the folded image is defined for all canonical coordinates with *N* (***u***; ***p***) ≥ 1. Requiring at least one valid sample is sufficient for folding because the purpose is to estimate the folded intensity from all available valid observations. The mask is transformed with the same geometric mapping *T_q_* as the image, which ensures that detector gaps, dead pixels, and saturated regions are excluded consistently during quadrant extraction, folding, registration, and loss evaluation.

The folded image can be interpreted as the best masked average estimate of the underlying symmetric diffraction intensity at coordinate ***u***. When the alignment is correct, corresponding valid quadrant samples contribute coherently to *Q̄*(***u***; ***p***). When the alignment is incorrect, samples from different quadrants may correspond to slightly different physical locations, producing blurring or reduced sharpness in the folded image. The quality of the folded image therefore depends directly on the accuracy of the center and orientation parameters.

### 3.5 Masked symmetry loss

A natural measure of alignment quality is the spread of corresponding quadrant intensities after masking invalid pixels. For each canonical folded coordinate ***u***, the local quadrant disagreement is the masked *standard deviation* across the valid quadrant samples,

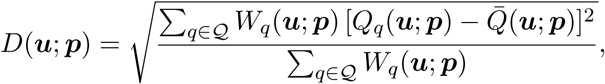

i.e. the population standard deviation (normalized by the valid-sample count *N* (***u***; ***p***) = ∑*_q_ W_q_*). This quantity is computed only when at least two valid quadrant samples are available, *N* (***u***; ***p***) ≥ 2. Coordinates with fewer than two valid samples are excluded because a single valid sample cannot provide a meaningful estimate of quadrant disagreement; this also prevents isolated valid samples from contributing misleading zero-dispersion values, so the loss measures disagreement among quadrants rather than the mere presence of valid data.

The global (unnormalized) symmetry loss is the sum of the local standard deviations over all qualifying coordinates,

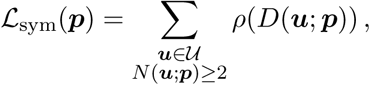

where *ρ*(·) is in general either the identity or a robust penalty such as the Huber loss. In this work, the realized loss takes *ρ* as the identity, so *L*_sym_ is exactly the summed per-pixel standard deviation. This loss directly measures how well the four reflected quadrants agree under the current estimate of the center and orientation. In a perfectly aligned and perfectly symmetric image, corresponding quadrant samples are identical and the loss is close to zero, up to image noise, interpolation effects, detector artifacts, and physical deviations from exact symmetry.

Because *L*_sym_ scales with image area and exposure, the score actually optimized and reported is the dimensionless *normalized* symmetry loss, obtained by dividing by the total foreground signal of the folded average,

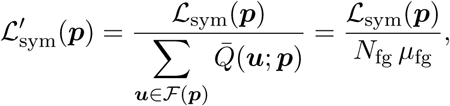

where the foreground set *F*(***p***) = {***u*** : *Q̄*(***u***; ***p***) *> τ*_Otsu_} collects the diffraction-signal coordinates of the folded-average image *Q̄*, the threshold *τ*_Otsu_ is set by Otsu’s method [13], which selects the threshold that maximizes between-class (foreground/background) variance, applied to the finite, positive values of *Q̄*. *N*_fg_ and *µ*_fg_ are the foreground pixel count (|*F* (***p***)|) and mean foreground intensity, respectively. Dividing by the total foreground signal ∑_***u****∈F Q̄*_ = *N*_fg_*µ*_fg_ makes *L^′^*_sym_ comparable across images of differing exposure and intensity scale.

The alignment-refinement problem is therefore ***p****^∗^* = arg min***_p_*** *L*_sym_(***p***). In practice the optimization is initialized from the rough alignment ***p***_0_ and estimates a small correction, Δ***p****^∗^* = arg min_Δ_***_p_*** *L*_sym_(***p***_0_ + Δ***p***), with refined alignment ***p****^∗^* = ***p***_0_ + Δ***p****^∗^*. When the rough alignment is expected to be close and large corrections are physically unlikely, a regularized version can be used,

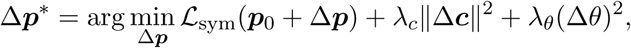

where the regularization terms discourage unrealistic deviations from the initial center and orientation.

Although *L*_sym_ is the most direct objective for folding quality, it can be difficult to optimize from a rough initial alignment: the loss is affected by interpolation, detector masks, sharp diffraction features, repeated structures, and changes in the valid-pixel set as the parameters vary. For this reason both direct optimization of *L*_sym_ and indirect methods, such as quadrant registration that provides a stable initialization for final local symmetry-loss refinement, are considered.

## 4 Methods

Four refinement strategies are considered: (1) hierarchical coarse-to-fine grid search on the masked symmetry loss; (2) ECC-based rigid registration; (3) local gradient-based refinement of the symmetry loss; and (4) a hybrid refinement approach that combines the previous methods.

### 4.1 Hierarchical coarse-to-fine grid search

A robust baseline is to perform a hierarchical coarse-to-fine grid search over the three correction parameters Δ***p*** = (Δ*c_x_,* Δ*c_y_,* Δ*θ*). Because the refinement problem has only three degrees of freedom, direct evaluation of the symmetry loss over a bounded parameter grid is computationally feasible. This approach avoids computing gradients and is therefore robust to interpolation artifacts, detector masks, sharp diffraction features, and non-smooth changes in the valid-pixel set. A coarse search over an initial grid G_1_ selects the candidate with the smallest loss; the search is then refined around that candidate using a smaller search window and smaller step sizes over successive levels G_2_*, . . .,* G*_L_*, and the final refined alignment is ***p****^∗^* = ***p***_0_ + Δ***p****_L_*. A representative three-level schedule is shown in Table 1.

**Table 1:** Example three-level coarse-to-fine grid-search schedule.

| Level | Center Range | Center Step | Angle Range | Angle Step |
| --- | --- | --- | --- | --- |
| 1 | $\pm 4$ px | 1.0 px | $\pm 2.0^\circ$ | $0.5^\circ$ |
| 2 | $\pm 1$ px | 0.25 px | $\pm 0.5^\circ$ | $0.1^\circ$ |
| 3 | $\pm 0.25$ px | 0.05 px | $\pm 0.1^\circ$ | $0.02^\circ$ |

For this schedule the number of evaluated candidates is (9×9×9)+(9×9×11) +(11×11×11) = 729 + 891 + 1331 = 2951, so the runtime is approximately 2951 *T*_eval_ where *T*_eval_ is the cost of one symmetry-loss evaluation. The method is therefore slower than local registration, but it remains practical for a three-parameter search, especially because candidate evaluations are independent and can be parallelized. Grid search also serves as a diagnostic: plotting the loss surface over (Δ*c_x_,* Δ*c_y_*) for selected Δ*θ* reveals whether the symmetry loss has a clear minimum near the rough alignment. If no clear minimum exists, the issue lies in the loss definition, mask handling, selected image regions, or assumed symmetry model rather than in the optimizer.

### 4.2 ECC registration and global correction fitting

A faster practical approach converts the symmetry-refinement problem into a set of image-registration problems. Given the current alignment ***p***_0_, the four quadrants are extracted into the common canonical coordinate system, *Q_q_*(***u***; ***p***_0_) = *I*(*T_q_*(***u***; ***p***_0_)). If the alignment is correct, all four quadrant images agree; if it is imperfect, they exhibit small relative translations and rotations. ECC-based rigid registration estimates a Euclidean transform *H_q_*(***u***) = *R*(*ϕ_q_*)***u*** +***t****_q_*, where *R*(*ϕ_q_*) is the rotation matrix and ***t****_q_* is the translation vector, that aligns a moving quadrant to a reference quadrant. The registration objective is correlation-based rather than a direct minimization of the four-quadrant folding loss, which makes ECC fast and effective when the rough alignment is reasonably close. The independently estimated quadrant transforms *H^∗^* must not be averaged directly. Because the quadrants are related by reflections, a true error in the global center and orientation induces different apparent transformations in different quadrants. The correct operation is to estimate the global correction Δ***p*** that best explains the observed quadrant-registration transforms. Using homogeneous coordinates, the mapping from canonical quadrant coordinates to original image coordinates under the rough alignment is

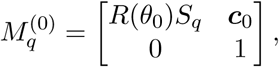

and for a refined alignment ***p***_1_ = ***p***_0_ + Δ***p*** the analogous matrix *M* ^(1)^ uses *R*(*θ*_0_ + Δ*θ*) and ***c***_0_ + Δ***c***. The change in coordinates induced by the refinement for quadrant *q* is *G_q_* (Δ***p***) = (*M_q_* ^(1)^)*^−^*^1^*M_q_* ^(0)^, so a candidate global refinement predicts a relative motion for every quadrant. The global correction is estimated by solving

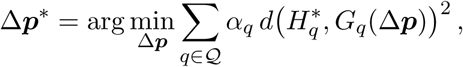

where *α_q_* is a confidence weight for quadrant *q* and *d*(·, ·) measures the disagreement between two rigid transforms, for example *d*(*H, G*)^2^ = *w_t_* ∥ ***t****_H_* − ***t****_G_* ∥ ^2^ + *w_θ_*(*ϕ_H_* − *ϕ_G_*)^2^. This projects the observed registrations onto the physically valid three-parameter alignment model rather than averaging transforms directly.

ECC was selected because it is fast, supports Euclidean motion, provides sub-pixel alignment, and is available in common image-processing libraries. Its main limitation is that it is a local registration method: it may fail or converge to an incorrect solution if the initial transform is too far from the correct alignment, if the image has weak structure, if the mask removes too much information, or if repeated layer-line patterns create ambiguous correlations. For this reason the ECC result is evaluated by the final folding objective *L*_sym_(***p***), not only by the pairwise registration score.

### 4.3 Local gradient-based refinement

The direct symmetry loss *L*_sym_(***p***) is the final objective of interest, but it is best optimized after a more stable method has produced a near-correct alignment. The difficulty is not that the objective is conceptually wrong but that its loss surface can be poorly behaved when the current estimate is not already close: interpolation of sharp layer lines and peaks makes the numerical gradient noisy; mask discontinuities change which pixels are valid as the parameters vary, so the objective is not smoothly varying; the three parameters are coupled, so the optimizer may reduce the loss by correcting the wrong combination of center and angle; repeated layer-line structures create ambiguous matches; and real patterns are only approximately symmetric, so a gradient optimizer may fit local physical asymmetries unless it is initialized close to the true alignment.

Direct gradient optimization is therefore treated as a local final refinement rather than a global search. Starting from an improved estimate ***p***_init_, the local refinement solves

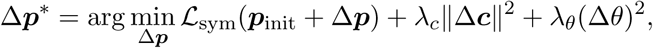

restricted to a small neighborhood |Δ*c_x_*|, |Δ*c_y_*| ≤ *r_c_*, |Δ*θ*| ≤ *r_θ_*. Because ***p***_init_ is already close to the optimum, the optimizer is less likely to be affected by large-scale non-convexity, ambiguous repeated structures, or incorrect quadrant correspondences, and it can correct small biases introduced by pairwise registration. In the implementation this stage uses L-BFGS-B.

### 4.4 Hybrid refinement pipeline

The candidate methods suggest a practical hybrid strategy. Hierarchical grid search is robust and useful as a diagnostic baseline but relatively expensive; direct gradient optimization targets the desired objective but is unreliable from a far initialization; ECC registration provides a faster intermediate solution by estimating relative quadrant motions that are then projected onto the physically valid three-parameter model. The simplest hybrid uses registration as the main correction mechanism, followed by a local refinement that directly minimizes the symmetry loss (referred to as “Reg+Grad”). In the evaluation this proved slightly suboptimal relative to brute-force search: across 56 image pairs from seven datasets, Reg+Grad differs on average ∼ 0.64 px and ∼ 0.3*^◦^* from where Search converges. Because symmetry evaluation is much cheaper on a cropped or downsampled image, we therefore append a third stage, a small-range local grid search on a cropped image, seeded from the Reg+Grad result (“Reg+Grad+Search”). The full procedure is summarized below; stopping after Stage 2 recovers the Reg+Grad variant.

1. **Initialization.** Set ***p*** ← ***p***_0_ (using MuscleX QF estimate for center and rotation), and record its masked symmetry loss *L*_sym_(***p***) as the current best.
2. **Stage 1 — iterative ECC registration (“Reg”).** Repeat for up to *K* passes (e.g. *K* = 5) or until convergence:

a. extract four quadrant images *Q_q_*(***u***; ***p***) and masks *W_q_*(***u***; ***p***) into common canonical frame;
b. choose the reference quadrant *r* as the one with the most valid pixels;
c. ECC-register each remaining quadrant *q* ≠ *r* to the reference, obtaining a rigid transform *H^∗^* and a confidence weight *w_q_*;
d. fit a single global correction Δ***p*** that best explains {*H^∗^*} under the reflection geometry, by weighted least squares;
e. update ***p*** ← ***p*** + Δ***p***; keep it as the new best if it lowers *L*_sym_; stop early when ∥ Δ***p*** ∥ is small or the loss stops decreasing.
3. **Stage 2 — local gradient refinement (yields “Reg+Grad”).** Minimize *L*_sym_ locally with L-BFGS-B in a small box |Δ***c***| ≤ *r_c_,* |Δ*θ*| ≤ *r_θ_* around ***p***; accept the result only if it reduces the loss, otherwise keep ***p***.
4. **Stage 3 — cropped local grid search (yields “Reg+Grad+Search”).** Crop the image and mask to a small window (e.g. 200 × 200 px) about the current center, and run a coarse-to-fine grid search over (Δ*c_x_,* Δ*c_y_,* Δ*θ*) on the cropped masked loss.
5. **Output.** Return the final alignment ***p****^∗^* ← ***p***.

Because the initial alignment is rough but close, only one or two registration passes are usually needed. Each stage after registration is accepted only if it lowers the masked symmetry loss, so the pipeline’s loss never increases.

### 4.5 Mask-aware ECC features and coarse-to-fine registration

#### ECC registration features

The registration routine exposes three optional, mask-aware features, all enabled by default. (1) *Mask erosion* shrinks the combined valid-pixel mask by a small elliptical structuring element (default 3 px) before registration, removing pixels near detector-gap boundaries where large intensity gradients can mislead the ECC optimizer, and re-calculates normalization and standard deviation for both quadrant images over the shared valid region. (2) *Re-seeding* warps the moving quadrant’s mask by the recovered transform after the first ECC pass, intersects it with the reference mask to obtain a post-registration overlap mask, and re-runs registration from the converged warp using this refined mask. (3) *Dynamic reference selection* chooses the quadrant with the most valid pixels as the ECC reference for each pair, rather than fixing the top-left quadrant, which is particularly useful on masked datasets.

#### Coarse-to-fine ECC registration

A two-stage registration runs ECC on a 1.5×-downsampled image for a few coarse passes, then uses the result to initialize full-resolution ECC for the remaining passes, as an alternative to single-scale registration.

### 4.6 Implementation considerations

Several implementation choices affect stability, accuracy, and runtime. The mappings *T_q_*(***u***; ***p***) generally sample the image at non-integer coordinates, so **interpolation** is required; bilinear interpolation is used, while the binary mask is sampled with nearest-neighbor interpolation or thresholded after interpolation so that partially invalid pixels are not treated as valid. For **mask handling**, the gap mask is transformed with the same geometry as the image, comparisons use only locations where both samples are valid, and the symmetry loss ignores coordinates with fewer than two valid samples.

## 5 Experimental Setup

### 5.1 Datasets

The evaluation uses real diffraction patterns from nine datasets spanning four tissue/preparation types, four of which carry a per-dataset detector-gap mask applied to every image (Table 2). The datasets differ in tissue (pig vs. mouse, cardiac vs. skeletal), preparation (skinned vs. intact), imaging medium (air vs. solution), and pattern strength (single frame vs. summed frames), giving a range of signal quality over which to test the refinement methods. The aggregate results in Section 6 are reported over the eight datasets that behave consistently under the reported and optimized scores; M10 is retained in the table for completeness but excluded from the aggregates as an outlier, for the reason discussed in Section 8.

**Table 2:** Datasets used in the evaluation. “Mask” indicates a per-dataset detector-gap mask applied to every image. “Images” is the number of frames scored per dataset. “Calib.” indicates whether a calibration image is available for the dataset.

| Dataset | Tissue / preparation | Images | Mask | Calib. | Notes |
| --- | --- | --- | --- | --- | --- |
| 2024.1213 | Skinned pig cardiac | 14 | no | yes<sup>†</sup> |  |
| 2025.0227 | Skinned pig cardiac | 5 | no | no |  |
| MPLL | Skinned pig cardiac | 5 | no | no | Clear layer lines |
| 2024.0508 | Skinned pig cardiac | 7 | yes | yes | High calcium |
| 2026.0409 | Skinned mouse cardiac | 5 | no | no | Strong pattern, shot in air |
| 2026.02.04 | Skinned mouse cardiac | 14 | no | yes |  |
| LP1 | Intact pig cardiac | 10 | yes | no | Single-frame (weak), in solution |
| LP3 | Intact pig cardiac | 10 | yes | yes | Single-frame (weak), in solution |
| M10 | Intact mouse skeletal | 10 | yes | no | Summed frames (strong); evaluation outlier |

<sup>†</sup>Calibration image present but unusable (see caption).

### 5.2 Baselines

The refinement methods are evaluated against three reference alignments. The primary one is the *MuscleX QF baseline*: the per-image center and rotation produced by the MuscleX quadrant-fold routine in the absence of a calibration image (Section 2), which is also the rough alignment ***p***_0_ from which every method is initialized. All methods are scored by how much they reduce the symmetry loss relative to this baseline.

For datasets with a calibration image, we add two stronger references. The *calibration baseline* applies a single per-dataset center — derived once from the AgBh calibration (Section 2) and shared by every image in the dataset — together with the MuscleX QF rotation. The *calibration-with-optimized-rotation* baseline keeps that calibration center but re-optimizes the rotation per image by grid search, isolating the value of an accurate, calibration-derived center from that of an accurate orientation. Comparing against these progressively stronger baselines quantifies how close the QF-initialized refinements come to a calibration-quality alignment (Section 6).

### 5.3 Metrics

#### Reported loss

Folding quality is measured by the normalized masked symmetry loss *L^′^_sym_*; lower is better. We distinguish the score that drives refinement from the score that is reported. Optimization, search, and registration are driven by a local score computed only on the specified crop size (Section 4), whereas all before/after values are reported with the whole-image score, which remains comparable across runs with different crop sizes and across the different refinement scripts.

#### Per-image and aggregate metrics

For each image, the loss and the mean residual intensity are recorded before and after refinement and summarized both as an absolute reduction Δ*L* and as a percentage Δ*L/L*_before_ × 100, and likewise for the residual. The magnitude of the applied correction is reported as the mean absolute center shift |Δ*C_x_*|, |Δ*C_y_*|, and rotation change |Δ*θ*|, together with the mean runtime.

### 5.4 Implementation details and reproducibility

#### Folding region

Unless noted, the folding objective is evaluated on a 600 × 600 px crop centered on the MuscleX QF center estimate. This window captures the main diffraction features, the meridional and equatorial reflections and the inner layer lines, while excluding peripheral noise and detector edges; fixing the crop keeps the loss comparable across methods and runs.

#### Optimizer settings

ECC registration uses 20 passes of 200 iterations each with Gaussian pre-blur *σ* = 1.0. The local gradient refinement runs up to 100 iterations with a maximum step of 1.0 px in center and 0.5*^◦^* in angle, and the cropped local search uses a 200 × 200 px window (Section 4).

#### Software

The pipeline runs on Python with MuscleX 2.0.0, NumPy 2.4, OpenCV 4.13, SciPy 1.17, and pyFAI 2026.5 / fabio 2025.10 for the calibration center audit. Per-image candidate evaluations are parallelized across CPU cores. The MuscleX quadrant-fold routine supplies the QF baseline and the symmetry-loss definition.

## 6 Results

### 6.1 Synthetic validation

A synthetic quadrant-symmetric image with a known center shift of 4.03 px and 2.0*^◦^* rotation, including noise and a gap mask, established the method ordering before the real-data evaluation. Grid search recovered near-zero parameter error but required 29.1 s; ECC registration reduced the center error to 0.125 px and the angular error to 0.0023*^◦^* in under 1 s; and ECC followed by gradient refinement produced the lowest final loss (2.145 × 10*^−^*^4^ vs. 2.146 × 10*^−^*^4^ for grid search) and the smallest center error (0.0065 px) at a total cost of 0.4 s. Direct gradient optimization from the naive alignment failed, consistent with the difficulty of the raw loss landscape from a rough start.

### 6.2 Multi-dataset evaluation

Across the eight datasets, refinement improves the symmetry loss and residual relative to the QF baseline for every method. Tables 3 and 4 aggregate the per-image metrics across all datasets in each group. On the unmasked datasets, which are all skinned cardiac muscle, the registration-based pipelines (Reg+Grad, Reg+Grad+Search, Reg only) match or exceed brute-force grid search at a fraction of its runtime. On the masked datasets, grid search retains the largest loss reduction, while the registration-based methods yield more modest gains. The masked datasets mostly consist of intact muscle, except for one skinned cardiac muscle dataset. The full per-dataset breakdown is reported in Appendix A, Tables 9 (unmasked) and 10 (masked).

**Table 3:** Unmasked datasets (5 datasets, 43 images): mean per-image improvement vs. QF baseline, aggregated across images.

| Method | $N$ | Loss reduction | | $\vert \Delta C_x\vert $ | $\vert \Delta C_y\vert $ | $\vert \Delta \theta\vert $ | Runtime |
| --- | --- | --- | --- | --- | --- | --- | --- |
| | | abs. | % | (px) | (px) | ($^\circ$) | |
| Search | 43 | 0.1986 | 13.99 | 0.75 | 0.68 | 1.06 | 425 |
| Reg+Grad | 43 | 0.2363 | 16.30 | 0.71 | 0.56 | 1.16 | 95 |
| Reg only | 43 | 0.2222 | 14.28 | 0.74 | 0.49 | 1.13 | 64 |
| Grad only | 43 | 0.0769 | 6.59 | 0.49 | 0.38 | 0.27 | 30 |
| Reg+Grad+Search | 43 | 0.2346 | 16.22 | 0.74 | 0.61 | 1.16 | 116 |
| Coarse-to-fine Reg | 43 | 0.2213 | 14.39 | 0.87 | 0.71 | 1.31 | 48 |

**Table 4:** Masked datasets (3 datasets, 27 images): mean per-image improvement vs. QF baseline, aggregated across images.

| Method | $N$ | Loss reduction | | $\vert \Delta C_x\vert $ | $\vert \Delta C_y\vert $ | $\vert \Delta \theta\vert $ | Runtime |
| --- | --- | --- | --- | --- | --- | --- | --- |
| | | abs. | % | (px) | (px) | ($^\circ$) | |
| Search | 27 | 0.1340 | 8.96 | 1.67 | 0.88 | 0.46 | 510 |
| Reg+Grad | 27 | 0.0409 | 2.91 | 1.43 | 0.98 | 0.36 | 98 |
| Reg only | 27 | 0.0151 | 1.14 | 1.41 | 0.99 | 0.34 | 61 |
| Grad only | 27 | 0.0279 | 1.82 | 0.35 | 0.28 | 0.13 | 39 |
| Reg+Grad+Search | 27 | 0.0849 | 5.62 | 1.50 | 0.85 | 0.43 | 122 |
| Coarse-to-fine Reg | 27 | 0.0119 | 0.98 | 1.43 | 0.99 | 0.36 | 55 |

Two control experiments isolate the cause of the lower registration improvement on masked data. First, datasets LP1 and LP3 were re-evaluated on images cropped to gap-free regions; registration then runs effectively without a mask. For the LP1 cropped version, Search reaches 5.80% (matching the full masked LP1) and Reg only reaches 1.04% (vs. 1.13% on the full set). For the LP3 cropped version, Search reaches 7.73% and Reg only 0.36%, both below the full masked LP3 (11.76% and 1.03%), indicating the cropped LP3 region carries less symmetry signal. Second, a synthetic “fake mask” was applied to the unmasked 2025 0227 dataset: every method’s loss reduction *improved* (Search 20.01 → 26.18%, Reg+Grad 23.13 → 31.07%), and the method ranking was unchanged. These controls, together with the different muscle types in the masked and unmasked datasets, indicate that poor registration improvement on masked datasets reflects dataset characteristics rather than a defect in mask handling.

### 6.3 Configuration studies

#### Registration feature ablation

Table 5 ablates the optional registration features against a plain registration baseline, aggregated across all datasets. Among mask erosion, re-seeding, dynamic reference selection, and their combinations, only enabling all features together clearly improves loss reduction. Registration with all features enabled is used as the “Reg” method in all other experiments.

**Table 5:** Registration feature ablation, combined across all masked and unmasked datasets (8 datasets): mean per-image improvement vs. QF baseline. Each row toggles one optional feature relative to the plain registration baseline; “All features” enables them together.

| Configuration | $N$ | Loss reduction | | Residual | $\vert \Delta C_x\vert $ | $\vert \Delta C_y\vert $ | $\vert \Delta \theta\vert $ | Runtime |
| --- | --- | --- | --- | --- | --- | --- | --- | --- |
| | | abs. | % | | (px) | (px) | ($^\circ$) | |
| None (baseline) | 70 | 0.0483 | 5.11 | 10.85 | 0.94 | 0.60 | 0.85 | 25 |
| Erode mask | 70 | 0.0480 | 5.06 | 10.88 | 0.95 | 0.61 | 0.87 | 26 |
| Reseed | 67 | 0.0416 | 4.63 | 9.85 | 0.96 | 0.62 | 0.81 | 42 |
| Erode + reseed | 70 | 0.0443 | 4.47 | 10.58 | 0.99 | 0.68 | 0.76 | 45 |
| Dynamic reference | 67 | 0.0422 | 4.59 | 9.74 | 1.01 | 0.69 | 0.80 | 24 |
| All features | 70 | 0.1412 | 9.14 | 27.28 | 1.00 | 0.68 | 0.83 | 63 |

#### Registration blur variations

Table 6 compares three levels of Gaussian pre-blur applied to the image before each ECC iteration (*σ* = 0, 1, 3), for the **Reg only** method, averaged across four datasets (LP1, 2025 0227, 2026 0409, 2026 02 04; *N* = 33 images). The three settings are nearly indistinguishable in aggregate. Per-dataset results show that no single value dominates across the individual datasets, which is why the default *σ* = 1 is used throughout the rest of the evaluations.

**Table 6:** ECC blur-sigma comparison for the Reg-only method, averaged across four datasets (LP1, 2025 0227, 2026 0409, 2026 02 04; 33 images): mean per-image improvement vs. QF baseline, weighted by image count.

| $\sigma$ | $N$ | Loss reduction | | Residual reduction | | $\vert \Delta C_x\vert $ | $\vert \Delta C_y\vert $ | $\vert \Delta \theta\vert $ | Runtime |
| --- | --- | --- | --- | --- | --- | --- | --- | --- | --- |
| | | abs. | % | abs. | % | (px) | (px) | ($^\circ$) | |
| 0 | 33 | 0.2599 | 15.87 | 0.9303 | 48.36 | 0.88 | 0.62 | 0.90 | 23 |
| 1 | 33 | 0.2629 | 16.19 | 1.0518 | 48.85 | 0.87 | 0.67 | 0.97 | 48 |
| 3 | 33 | 0.2648 | 16.51 | 1.0932 | 48.77 | 0.85 | 0.61 | 0.99 | 19 |

#### Crop size variations

Three folding-region configurations are compared, with results averaged across four datasets (2025 0227, 2026 0409, MPLL, LP3):

- **600**×**600**: *r_x_* = 600, *r_y_*= 600 (default, used in the rest of the evaluations).
- **200**×**200**: *r_x_* = 200, *r_y_*= 200 (a smaller region where most of the intensity information is retained).
- **100**×**600**: *r_x_* = 600, *r_y_*= 100 (an elongated “equator region”).

Five methods are evaluated under each crop configuration (Search, Reg+Grad, Reg only, Grad only, Coarse-to-fine), all vs. the QF baseline. Crop size matters in a strongly method-dependent way, so the large default is not uniformly best (Table 7):

- **Coarse-to-fine** is by far the most crop-sensitive: it needs the full 600 × 600 and collapses at the smaller crops due to the downsampling, where most information is lost.
- **Search** peaks at 600 × 600, but the two smaller crops come within a percentage point at a fraction of the runtime.
- **Reg+Grad** and **Reg only** are marginally better at 200 × 200 than at the default, and **Grad only** improves with smaller cropping regions.

**Table 7:** Crop-size comparison, averaged across four datasets (2025 0227, 2026 0409, MPLL, LP3; 24 images): mean per-image improvement vs. QF baseline. Bold shows the best loss reduction crop per method.

| Method | Crop | $N$ | Loss reduction | | Residual<br>red. % | Runtime<br>(s) |
| --- | --- | --- | --- | --- | --- | --- |
|  |  |  | abs. | % |  |  |
| Search | 600×600 | 24 | 0.1335 | **11.88** | 19.41 | 625 |
| Search | 200×200 | 24 | 0.1162 | 10.88 | 27.69 | 19 |
| Search | 100×600 | 24 | 0.1148 | 10.82 | 27.38 | 28 |
| Reg+Grad | 600×600 | 24 | 0.0654 | 8.83 | 16.92 | 182 |
| Reg+Grad | 200×200 | 24 | 0.0813 | **10.06** | 27.59 | 12 |
| Reg+Grad | 100×600 | 24 | 0.0624 | 8.21 | 25.57 | 16 |
| Reg only | 600×600 | 24 | 0.0389 | 6.16 | 12.35 | 122 |
| Reg only | 200×200 | 24 | 0.0387 | **6.49** | 19.21 | 10 |
| Reg only | 100×600 | 24 | 0.0292 | 5.28 | 17.94 | 14 |
| Grad only | 600×600 | 24 | 0.0433 | 5.27 | 9.74 | 48 |
| Grad only | 200×200 | 24 | 0.0603 | 6.29 | 17.46 | 1 |
| Grad only | 100×600 | 24 | 0.0650 | **6.38** | 16.78 | 2 |
| Coarse-to-fine | 600×600 | 24 | 0.0417 | **7.08** | 3.12 | 76 |
| Coarse-to-fine | 200×200 | 24 | 0.0033 | 0.47 | 3.32 | 7 |
| Coarse-to-fine | 100×600 | 24 | 0.0022 | 0.44 | 3.00 | 9 |

The smaller crops also cut runtime by one to two orders of magnitude.

### 6.4 Comparison with a calibration-image baseline

All methods in this experiment are initialized from the per-image **MuscleX QF estimates** and evaluated against two reference baselines: (1) the **MuscleX QF baseline** (per-image auto QF center and rotation), and (2) a **calibration baseline** derived from a single calibration image whose center serves as the fixed reference for the entire dataset. The calibration baseline is considered in two variants: one using the MuscleX QF rotation, the other re-optimizing rotation by grid search.

Table 8 aggregates the per-image metrics across the three datasets with a usable calibration image (2024 0508, 2026 02 04, LP3; 31 images). The table is split into three panels: improvement relative to the **MuscleX QF baseline** (upper), relative to the **calibration baseline** with MuscleX QF rotation (middle), and relative to the **calibration baseline with optimized rotation** (lower). A negative entry against a calibration baseline means the calibration center/rotation is already at least as good as the refined solution. This confirms that a calibration image is effectively the optimal reference when available; without it, Search (or hybrid registration + local search) is the next best option.^1^ The full per-dataset breakdown is given in Appendix B (Table 11).

**Table 8:** Calibration comparison, aggregated across the three datasets with a usable calibration image (2024 0508, 2026 02 04, LP3; 31 images): mean per-image improvement weighted by image count, relative to the QF baseline (upper), the calibration baseline (middle), and the calibration baseline with optimized rotation (lower). All methods are initialized from QF estimates. Negative values indicate that the method performs worse than the respective baseline, indicating that calibration with the optimized rotation is the best option. When a calibration image is not available, Search is the next best option.

| Method | $N$ | Loss Reduction | | $\vert \Delta C_x\vert $ (px) | $\vert \Delta C_y\vert $ (px) | $\vert \Delta \theta\vert $ (°) | Runtime (s) |
| --- | --- | --- | --- | --- | --- | --- | --- |
|  |  | abs. | % |  |  |  |  |
| *vs. QF baseline* |  |  |  |  |  |  |  |
| Search | 31 | 0.2899 | 15.21 | 1.42 | 0.82 | 0.74 | 452 |
| Reg+Grad | 31 | 0.2799 | 14.48 | 1.27 | 0.83 | 0.74 | 85 |
| Grad only | 31 | 0.0815 | 4.31 | 0.32 | 0.22 | 0.13 | 37 |
| Reg only | 31 | 0.2646 | 13.57 | 1.25 | 0.78 | 0.72 | 50 |
| *vs. Calibration baseline* |  |  |  |  |  |  |  |
| Search | 31 | 0.3850 | 18.54 | 0.37 | 0.25 | 0.71 | 452 |
| Reg+Grad | 31 | 0.3749 | 17.53 | 0.36 | 0.73 | 0.67 | 85 |
| Grad only | 31 | 0.1765 | 8.10 | 1.18 | 0.61 | 0.74 | 37 |
| Reg only | 31 | 0.3597 | 16.52 | 0.37 | 0.78 | 0.66 | 50 |
| *vs. Calibration baseline with optimized rotation* |  |  |  |  |  |  |  |
| Search | 31 | -0.0242 | -1.73 | 0.37 | 0.25 | 0.27 | 452 |
| Reg+Grad | 31 | -0.0343 | -2.45 | 0.36 | 0.73 | 0.39 | 85 |
| Grad only | 31 | -0.2330 | -16.70 | 1.18 | 0.61 | 0.85 | 37 |
| Reg only | 31 | -0.0496 | -3.48 | 0.37 | 0.78 | 0.41 | 50 |

Figure 2 shows the alignments visually on a representative LP3 image. The raw pattern is off-center and partly occluded by detector gaps; the AgBh calibrant exposure (from which the calibration center is derived) shows the powder ring used for the fit; and the QF and calibration-with-optimized-rotation alignments both recenter the pattern, with the calibration variant providing the reference against which the QF-initialized refinements are compared.

**Figure 2:**
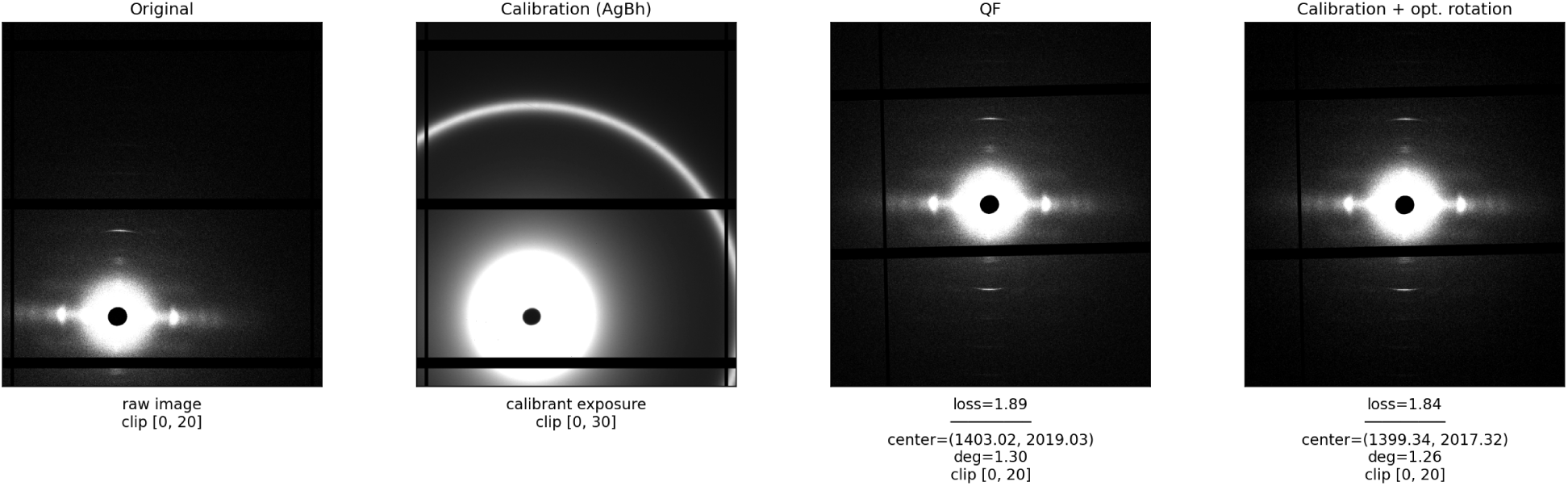
Calibration comparison on a representative LP3 image. From left to right: the original image; the AgBh calibrant exposure used to derive the calibration center; the image aligned with the MuscleX QF estimate (loss and center/rotation shown); and the image aligned with the calibration center and a per-image optimized rotation. Intensities are clipped per panel (the weak tissue pattern and the bright calibrant ring use different scales).

#### Meridional projection examples (LP3)

Each panel in Figures 3 and 4 shows the vertical summed-intensity projection within a 20 px-wide band centered on the diffraction meridian (*c_x_* = 0 in the refined coordinate frame) for two representative LP3 images. Columns, left to right: **Search**, **Registration**, **Registration-and-Gradient**, **Calibrated**, **MuscleX QF Auto**, **Original**. The Calibration column uses the MuscleX center calibration with optimized rotation (grid search). All panels, except the *Original* column, show projections from quadrant folded images. The *Original* column is the original image translated and rotated to the Search solution coordinates, so it serves as the reference against which the other methods are compared; a method whose profile matches the Original most closely on the main high-angle features has found the best center and rotation that, when folded, best preserves the original structure. Search and Calibration consistently best reproduce the Original features (peak positions and relative intensities); Registration/Reg+Grad and QF show some deviations. For image 105, both Calibration and Search preserve M6 (≈ −800 in the *y* coordinate); for image 115, the Calibration projection is less accurate on M6 and its rotation angle is the most different from the rest.

**Figure 3:**
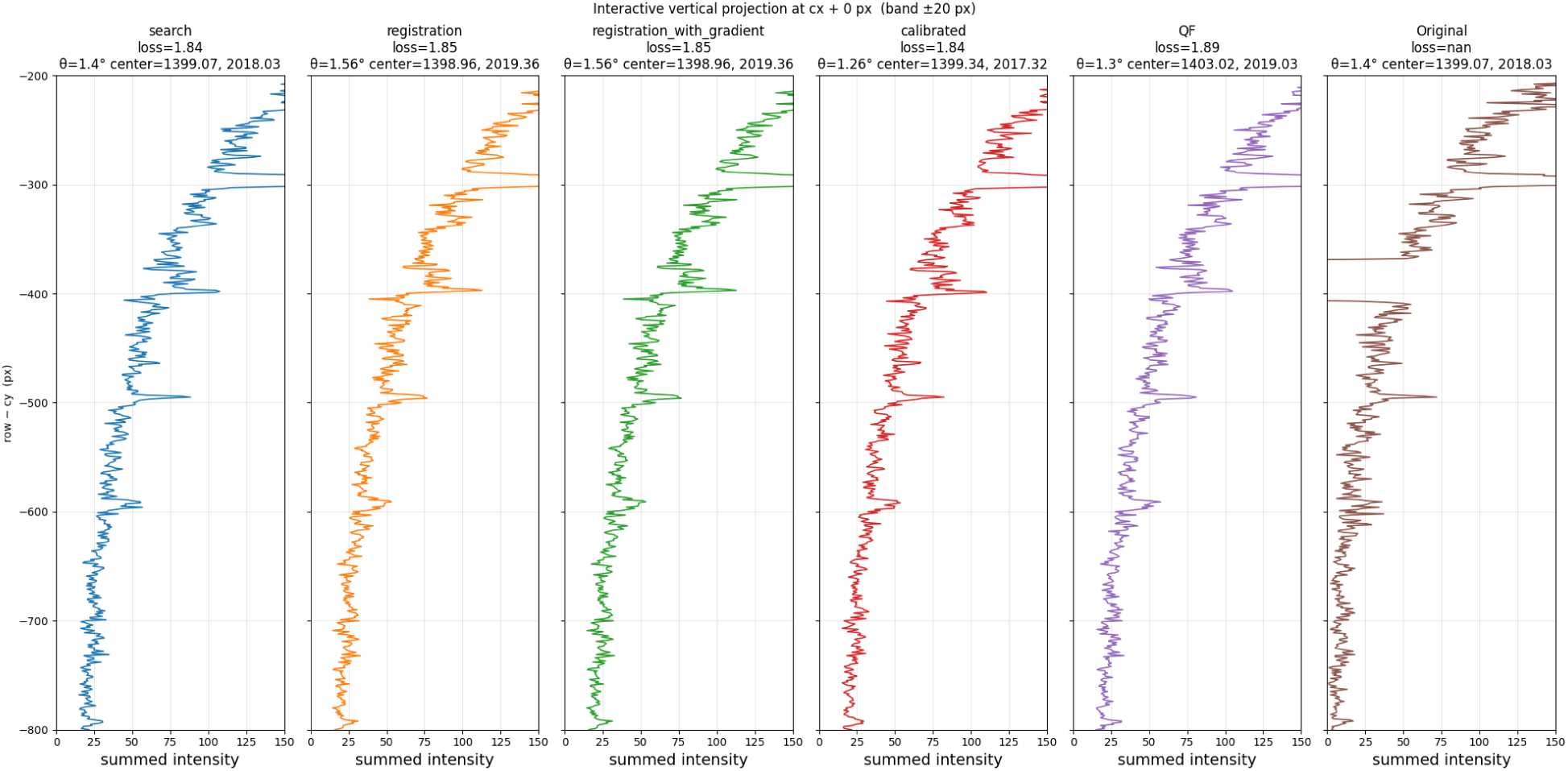
LP3, image 105: meridional projections (20 px band, *c_x_* = 0). Search and Calibration closely reproduce the Original profile.

**Figure 4:**
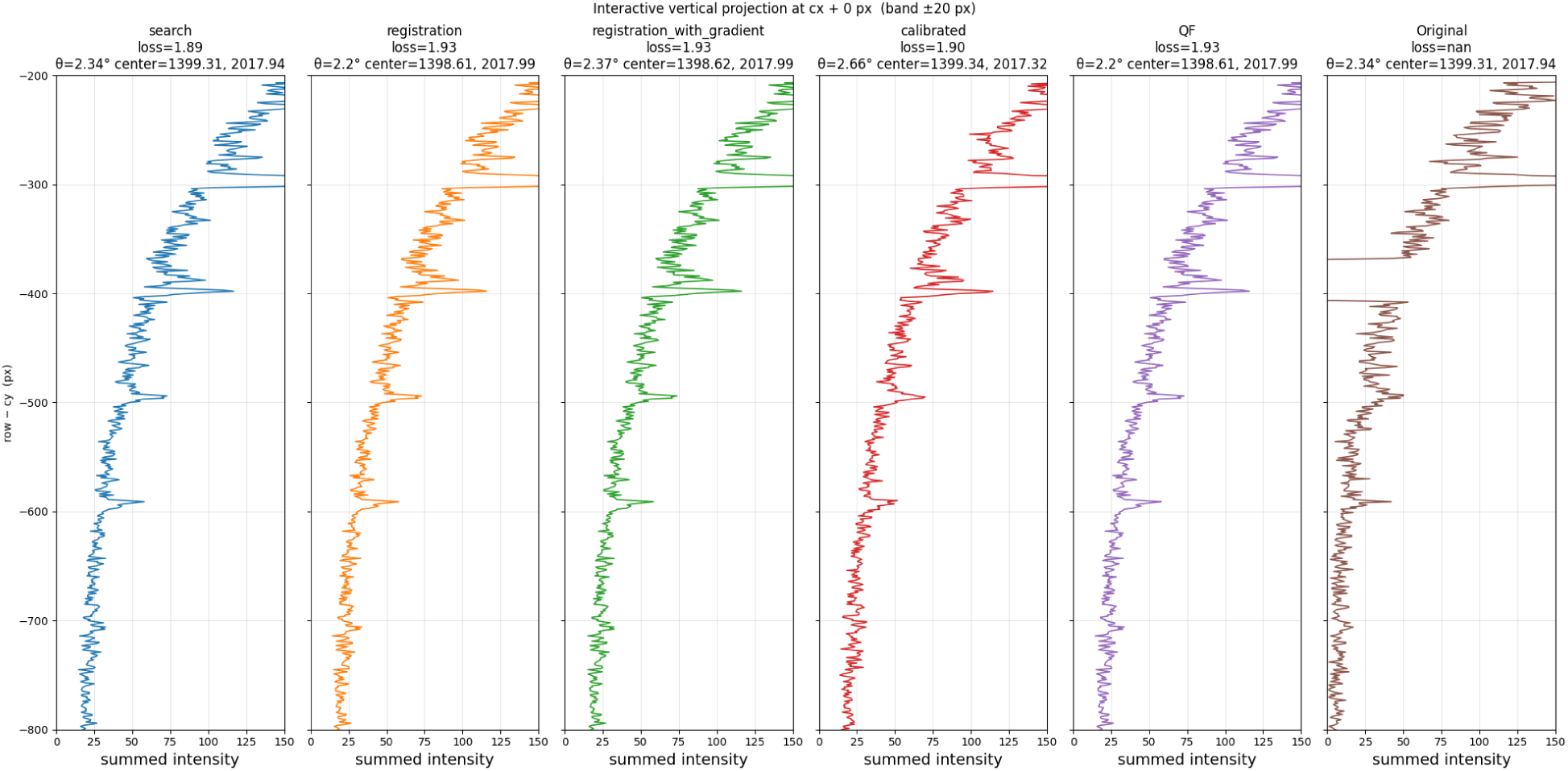
LP3, image 115: meridional projections (20 px band, *c_x_* = 0). Search, Calibration and Reg+Grad closely reproduce the Original profile.

### 6.5 Comparison of MuscleX calibration-derived center to pyFAI

We additionally compared pyFAI calibration against the MuscleX QF reference. Based on limited testing, we conclude that in pyFAI the reliability of the recovered beam center depends on the calibration image quality and peak-finder choice. In pyFAI, the detector-sample geometry is determined as a set of PONI parameters from which the beam center is derived, using a powder calibrant (AgBh) that produces rings at known radii. Ideally, the calibration image should have multiple well-defined rings for pyFAI geometry evaluation. From one user-clicked seed per ring, a peak-finder collects ring pixels and GeometryRefinement minimizes the chi-squared residual between observed and predicted ring positions over (dist, poni1, poni2, rot1, rot2). There are two main algorithms pyFAI uses for peak finding on the rings: Massif (default) and Watershed. Massif flood-fills a connected basin from each seed, so it can miss arcs broken by gaps or low intensity, and its random peak sampling introduces run-to-run variance. Watershed pre-segments the whole image and collects all labeled peaks in an annular mask, giving deterministic global coverage that depends on the seed only for the radius.

When using the Massif peak-finder, pyFAI recovers the center to sub-pixel agreement with the MuscleX QF reference on three of four test images but fails on LP3, where its column estimate scatters by ±4.3 px across runs and results in ∼ 4 px difference from the reference. When using the Watershed peak-finder, the result is fully deterministic (std = 0 on every parameter) yet converges to a wrong center on the two images with gaps or a single usable ring, missing the true center. Each method was run five times per image.

On the clean Pilatus1M reference (three sharp rings), both methods agree within ∼ 0.15 px. On 2024 0508 (two rings, no gaps), all three estimates agree within ∼ 0.2 col / ∼ 0.6 row px. On 2024 1213 (faint, single usable ring), Massif agrees with MuscleX to within ∼ 0.6 col / 1 row px, while Watershed places the center with row and column transposed — a local minimum produced by the near-diagonal symmetry of a single centered ring on a square detector. On LP3 (gaps, two rings), Massif’s converged center is ∼ 4 px from MuscleX.

For reliable use, Massif with at least two gap-free, well-defined rings is the configuration most likely to align with the MuscleX baseline. The MuscleX calibration only calculates the center coordinates and the radius, rather than the full set of PONI parameters; it has low run-to-run variance and provides a reliable reference for comparison.

## 7 Discussion

The evaluation supports a consistent picture of how the refinement strategies relate to one another.

### Registration recovers most of the achievable improvement at a fraction of the cost

On the unmasked datasets, the registration-based pipelines run four to seven times faster than brute-force search (Table 3) while matching or exceeding its loss reduction, and the added gradient step closes most of the remaining gap to Search.

### The speed/accuracy trade-off is method- and crop-dependent

Search is the most accurate but by far the slowest, and most of its accuracy survives at much smaller crops (Table 7). The hybrid Reg+Grad+Search variant is the intended default when both accuracy and speed matter: it inherits the speed of registration and recovers the last increment of accuracy with a cheaper cropped local search.

### Lower symmetry loss corresponds to physically better alignment

The loss is a proxy for folding quality, so it is important that the ranking it induces agrees with an independent physical check. The meridional-projection comparison on LP3 (Figures 3 and 4) shows that the methods that achieve the lowest loss also reproduce the original high-angle peak positions and relative intensities most faithfully.

### A calibration image is the effective upper bound

When a usable calibration image is available, applying its center together with a per-image optimized rotation matches or exceeds every QF-initialized refinement. The refinement methods are best understood as recovering a calibration-quality alignment. This also explains why the gains separate into a center contribution and a rotation contribution: against the calibration center with the QF rotation, the refinements still improve substantially, but against the calibration center with optimized rotation, they do not, indicating that an accurate center is the harder part to recover and the rotation is comparatively easy to optimize once the center is fixed.

## 8 Limitations

Several limitations qualify the results and indicate where the method should be applied with care.

### Local-versus-whole-image scoring can disagree

Refinement is driven by a local score computed on a crop, whereas the reported before/after values use the whole-image score. The two usually agree, but not always. This mismatch is the reason why the M10 dataset behaves as an outlier. M10 is therefore excluded from the aggregates rather than allowed to distort them, and the discrepancy is a property of the scoring procedure, not of the optimizer.

### Some dataset types yield smaller registration gains

Intact masked datasets seem to produce the lowest gains with registration. Local search in the “Reg+Grad+Search” approach helps to close this gap, but further analysis is needed to understand the causes of the discrepancy.

### The calibration baseline rests on a single calibration image per dataset

That center is assumed valid for every frame, so per-frame detector shift is not captured. The calibration path also has a hard failure mode: the 2024 1213 dataset had to be dropped from the calibration comparison because its calibration ring extended beyond the detector boundary, which the MuscleX calibration procedure does not handle at this time.

### The pyFAI reliability study is small and self-referential

It covers four images and treats the MuscleX QF center as the reference, so it measures agreement between two calibration routes rather than absolute accuracy against an independently known center; a larger, ground-truthed study would be needed to draw quantitative conclusions about the quality of center finding.

## 9 Conclusion and Recommendations

We formulated masked quadrant folding as the minimization of a normalized, mask-aware four-quadrant symmetry loss, and evaluated a family of refinement strategies that correct a rough per-image alignment against this objective. Direct gradient optimization from the rough alignment was found to be unreliable; brute-force grid search is robust and interpretable but slow; and ECC-based quadrant registration, projected onto a single physically valid center/orientation correction, provides a fast near-correct alignment. Combining the approaches — registration, then a local gradient step, then a cheap cropped local search (Reg+Grad+Search) — recovers the accuracy of brute-force search with lower runtime. When a calibration image is available, applying its center with a per-image optimized rotation is effectively optimal.

### Recommendations

- *Routine refinement without a calibration image:* use registration with all mask-aware features enabled, followed by local gradient refinement and a small cropped local search (Reg → Grad → 200 × 200 search). This gives most of the achievable loss reduction at low cost.
- *Reference / diagnostic:* use Search at a larger cropping region when an accuracy ceiling or a loss-surface check is needed, accepting its higher runtime.
- *When a calibration image is available:* adopt the calibration center with a per-image optimized rotation as the alignment.

These approaches are implemented in the MuscleX QF module starting with version 2.0 [11].

## A Per-dataset result tables

The per-dataset breakdown is split by group: Table 9 covers the five unmasked datasets and Table 10 the three masked datasets, all scored against the QF baseline.

**Table 9:**
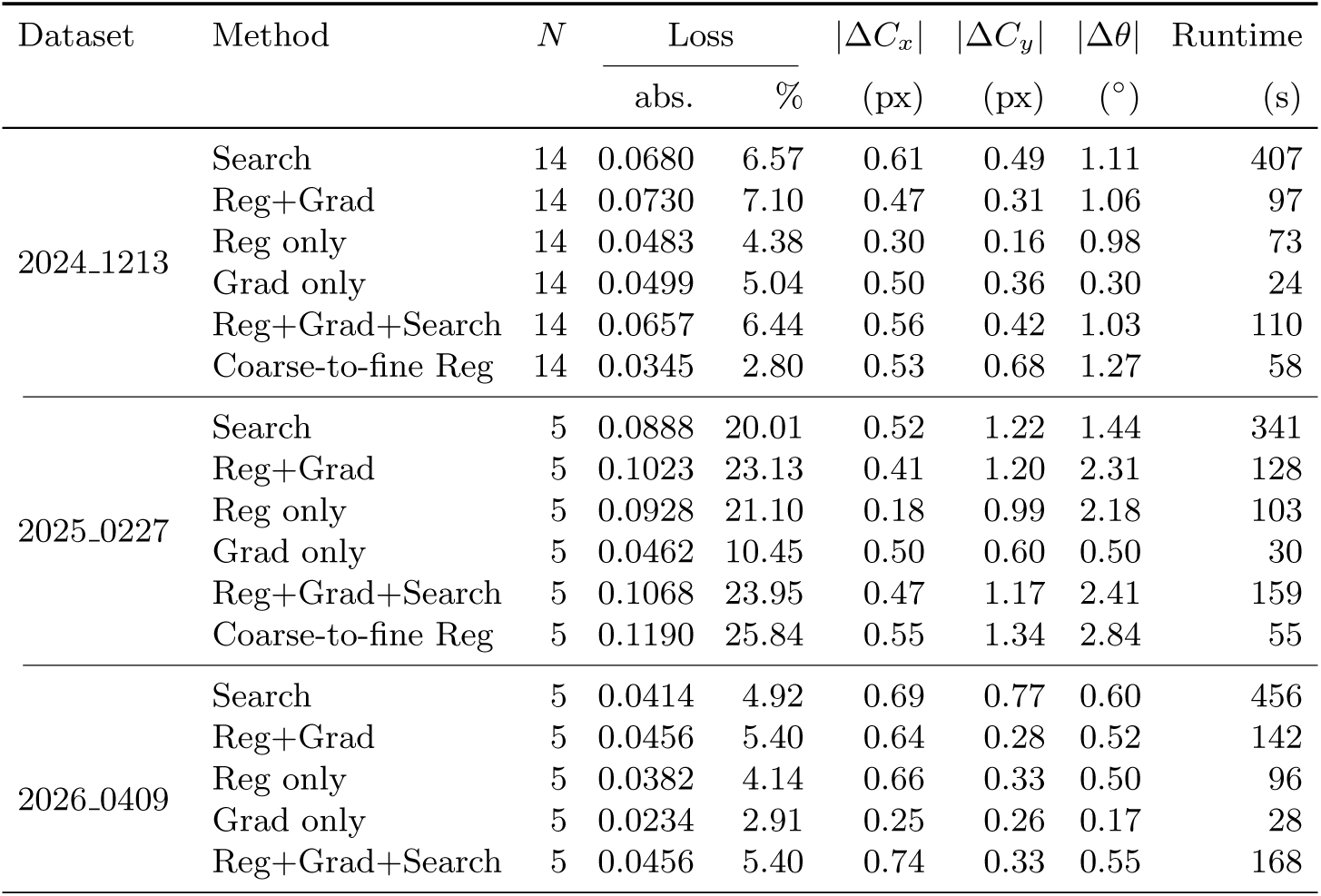
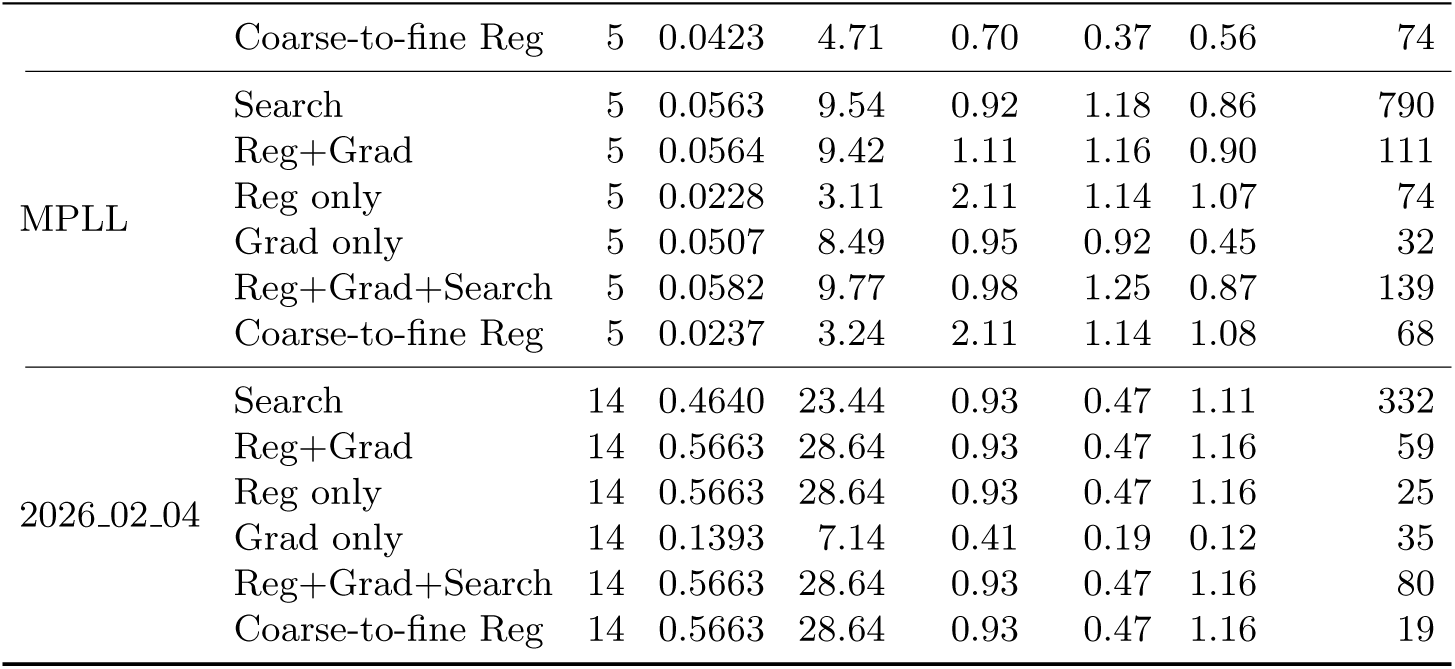
Unmasked datasets: per-dataset mean per-image improvement vs. QF baseline, by method. “Loss” gives the absolute and percentage reduction; |Δ*C_x_*|, |Δ*C_y_*| are center shifts (px), |Δ*θ*| the angle change (*^◦^*), and Runtime is in seconds.

**Table 10:**
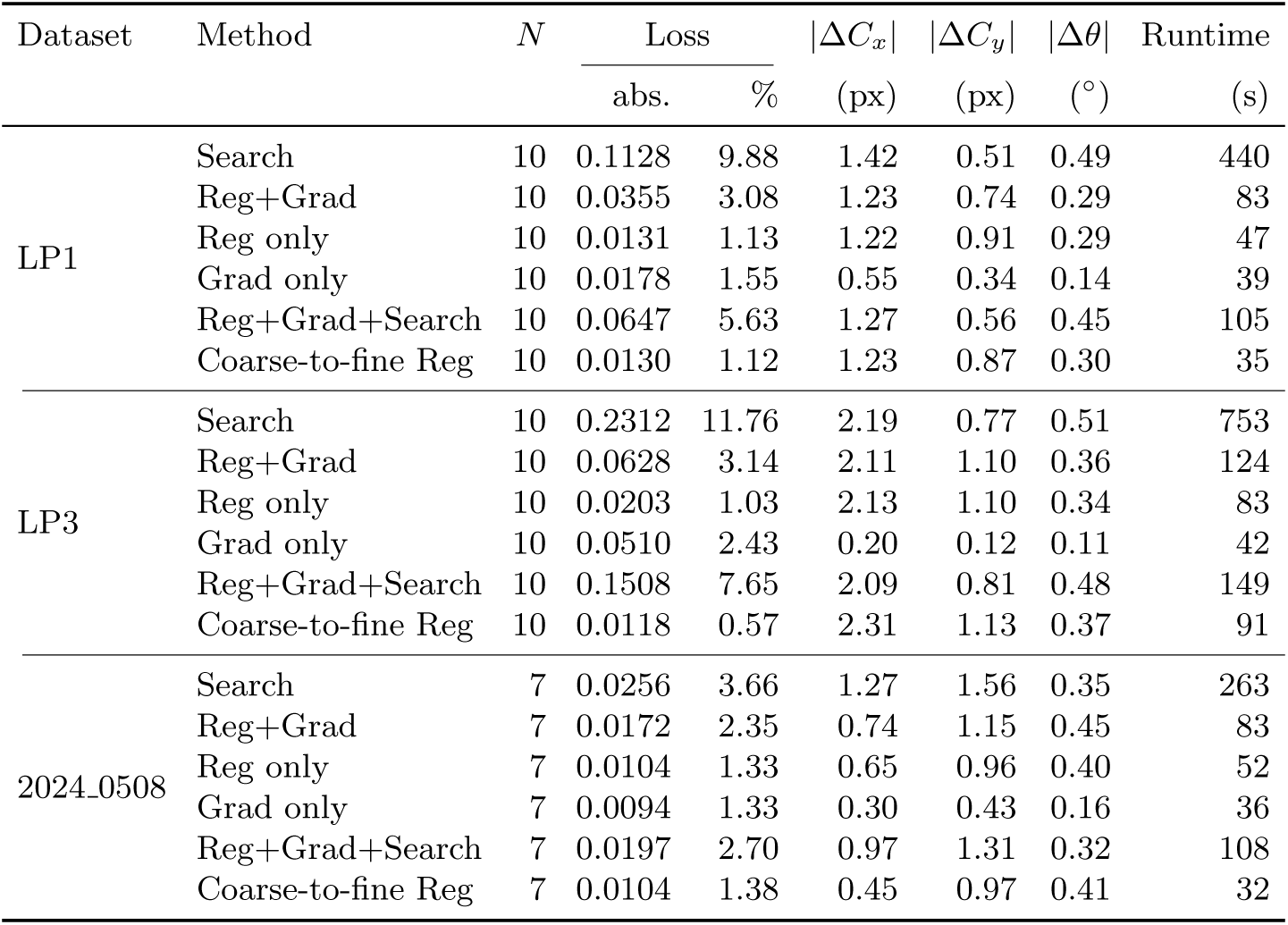
Masked datasets: per-dataset mean per-image improvement vs. QF baseline, by method. Columns as in Table 9.

| Dataset | Method | $N$ | Loss | | $\vert \Delta C_x\vert $ | $\vert \Delta C_y\vert $ | $\vert \Delta \theta\vert $ | Runtime |
| --- | --- | --- | --- | --- | --- | --- | --- | --- |
| | | | abs. | % | (px) | (px) | ($^\circ$) | |
| LP1 | Search | 10 | 0.1128 | 9.88 | 1.42 | 0.51 | 0.49 | 440 |
|  | Reg+Grad | 10 | 0.0355 | 3.08 | 1.23 | 0.74 | 0.29 | 83 |
|  | Reg only | 10 | 0.0131 | 1.13 | 1.22 | 0.91 | 0.29 | 47 |
|  | Grad only | 10 | 0.0178 | 1.55 | 0.55 | 0.34 | 0.14 | 39 |
|  | Reg+Grad+Search | 10 | 0.0647 | 5.63 | 1.27 | 0.56 | 0.45 | 105 |
|  | Coarse-to-fine Reg | 10 | 0.0130 | 1.12 | 1.23 | 0.87 | 0.30 | 35 |
| LP3 | Search | 10 | 0.2312 | 11.76 | 2.19 | 0.77 | 0.51 | 753 |
|  | Reg+Grad | 10 | 0.0628 | 3.14 | 2.11 | 1.10 | 0.36 | 124 |
|  | Reg only | 10 | 0.0203 | 1.03 | 2.13 | 1.10 | 0.34 | 83 |
|  | Grad only | 10 | 0.0510 | 2.43 | 0.20 | 0.12 | 0.11 | 42 |
|  | Reg+Grad+Search | 10 | 0.1508 | 7.65 | 2.09 | 0.81 | 0.48 | 149 |
|  | Coarse-to-fine Reg | 10 | 0.0118 | 0.57 | 2.31 | 1.13 | 0.37 | 91 |
| 2024.0508 | Search | 7 | 0.0256 | 3.66 | 1.27 | 1.56 | 0.35 | 263 |
|  | Reg+Grad | 7 | 0.0172 | 2.35 | 0.74 | 1.15 | 0.45 | 83 |
|  | Reg only | 7 | 0.0104 | 1.33 | 0.65 | 0.96 | 0.40 | 52 |
|  | Grad only | 7 | 0.0094 | 1.33 | 0.30 | 0.43 | 0.16 | 36 |
|  | Reg+Grad+Search | 7 | 0.0197 | 2.70 | 0.97 | 1.31 | 0.32 | 108 |
|  | Coarse-to-fine Reg | 7 | 0.0104 | 1.38 | 0.45 | 0.97 | 0.41 | 32 |

## B Per-dataset calibration tables

Table 11 gives the full per-dataset breakdown underlying the aggregate in Table 8. For each dataset the rows are grouped into three panels: improvement relative to the **QF baseline**, relative to the **calibration baseline** (MuscleX QF rotation), and relative to the **calibration baseline with optimized rotation**. All methods are initialized from the QF estimates. A negative loss-reduction entry against a calibration baseline indicates that the calibration center/rotation is already at least as good as the refined solution.

**Table 11:** Per-dataset calibration comparison: mean per-image metrics relative to the QF baseline, the calibration baseline, and the calibration baseline with optimized rotation, all initialized from QF estimates. “Loss abs.” and “Loss %” give the absolute and percentage loss reduction; |Δ*C_x_*|, |Δ*C_y_*| are center shifts (px), |Δ*θ*| the angle change (*^◦^*), and Runtime is in seconds.

| Dataset | Baseline | Method | $N$ | Loss | | $\vert \Delta C_x\vert $ | $\vert \Delta C_y\vert $ | $\vert \Delta\theta\vert $ | Runtime |
| --- | --- | --- | --- | --- | --- | --- | --- | --- | --- |
| | | | | abs. | % | (px) | (px) | ($^\circ$) | (s) |
| 2024.0508 | QF | Search | 7 | 0.0256 | 3.66 | 1.27 | 1.56 | 0.35 | 263 |
|  |  | Reg+Grad | 7 | 0.0172 | 2.35 | 0.74 | 1.15 | 0.45 | 83 |
|  |  | Grad only | 7 | 0.0094 | 1.33 | 0.30 | 0.43 | 0.16 | 36 |
|  |  | Reg only | 7 | 0.0104 | 1.33 | 0.65 | 0.96 | 0.40 | 52 |
|  | Calib. | Search | 7 | 0.0197 | 2.92 | 1.26 | 0.41 | 0.23 | 263 |
|  |  | Reg+Grad | 7 | 0.0113 | 1.59 | 0.96 | 0.54 | 0.47 | 83 |
|  |  | Grad only | 7 | 0.0035 | 0.54 | 1.21 | 0.85 | 0.42 | 36 |
|  |  | Reg only | 7 | 0.0045 | 0.60 | 1.06 | 0.72 | 0.42 | 52 |
|  | Calib.+rot | Search | 7 | -0.0011 | -0.25 | 1.26 | 0.41 | 0.39 | 263 |
|  |  | Reg+Grad | 7 | -0.0096 | -1.69 | 0.96 | 0.54 | 0.54 | 83 |
|  |  | Grad only | 7 | -0.0173 | -2.73 | 1.21 | 0.85 | 0.48 | 36 |
|  |  | Reg only | 7 | -0.0164 | -2.76 | 1.06 | 0.72 | 0.61 | 52 |
| 2026.02.04 | QF | Search | 14 | 0.4640 | 23.44 | 0.93 | 0.48 | 1.11 | 332 |
|  |  | Reg+Grad | 14 | 0.5663 | 28.64 | 0.93 | 0.47 | 1.16 | 59 |
|  |  | Grad only | 14 | 0.1393 | 7.14 | 0.41 | 0.19 | 0.12 | 35 |
|  |  | Reg only | 14 | 0.5663 | 28.64 | 0.93 | 0.47 | 1.16 | 25 |
|  | Calib. | Search | 14 | 0.7118 | 32.78 | 0.01 | 0.01 | 1.14 | 332 |
|  |  | Reg+Grad | 14 | 0.8140 | 37.41 | 0.00 | 0.00 | 1.09 | 59 |
|  |  | Grad only | 14 | 0.3871 | 17.82 | 0.52 | 0.28 | 1.16 | 35 |
|  |  | Reg only | 14 | 0.8140 | 37.41 | 0.00 | 0.00 | 1.09 | 25 |
|  | Calib.+rot | Search | 14 | -0.0533 | -3.85 | 0.01 | 0.01 | 0.07 | 332 |
|  |  | Reg+Grad | 14 | 0.0489 | 2.44 | 0.00 | 0.00 | 0.02 | 59 |
|  |  | Grad only | 14 | -0.3780 | -28.05 | 0.52 | 0.28 | 1.06 | 35 |
|  |  | Reg only | 14 | 0.0489 | 2.44 | 0.00 | 0.00 | 0.02 | 25 |
| LP3 | QF | Search | 10 | 0.2312 | 11.76 | 2.20 | 0.77 | 0.51 | 753 |
|  |  | Reg+Grad | 10 | 0.0628 | 3.14 | 2.11 | 1.10 | 0.36 | 124 |
|  |  | Grad only | 10 | 0.0510 | 2.43 | 0.20 | 0.12 | 0.11 | 42 |
|  |  | Reg only | 10 | 0.0203 | 1.03 | 2.13 | 1.10 | 0.34 | 83 |
|  | Calib. | Search | 10 | 0.1831 | 9.55 | 0.26 | 0.46 | 0.45 | 753 |
|  |  | Reg+Grad | 10 | 0.0147 | 0.85 | 0.45 | 1.90 | 0.22 | 124 |
|  |  | Grad only | 10 | 0.0029 | -0.22 | 2.07 | 0.90 | 0.38 | 42 |
|  |  | Reg only | 10 | -0.0278 | -1.58 | 0.42 | 1.92 | 0.24 | 83 |
|  | Calib.+rot | Search | 10 | 0.0003 | 0.19 | 0.26 | 0.46 | 0.46 | 753 |
|  |  | Reg+Grad | 10 | -0.1681 | -9.82 | 0.45 | 1.90 | 0.80 | 124 |
|  |  | Grad only | 10 | -0.1799 | -10.60 | 2.07 | 0.90 | 0.83 | 42 |
|  |  | Reg only | 10 | -0.2106 | -12.27 | 0.42 | 1.92 | 0.82 | 83 |

## Footnotes

1 A fourth dataset with calibration data, 2024 1213, is excluded from this comparison: its calibration ring extends beyond the detector boundary, a case the MuscleX calibration procedure does not currently handle, so no reliable calibration center can be derived for it.

